# Tissue-Specific Metabolomic Reprogramming Determines the Disease Pathophysiology of Sars-Cov-2 Variants in Hamster Model

**DOI:** 10.1101/2024.02.25.581989

**Authors:** Urvinder Kaur Sardarni, Anoop T Ambikan, Arpan Acharya, Samuel D Johnson, Sean N. Avedissian, Akos Vegvan, Ujjwal Neogi, Siddappa N. Byrareddy

## Abstract

Despite significant effort, a clear understanding of host tissue-specific responses and their implications for immunopathogenicity against the severe acute respiratory syndrome coronavirus 2 (SARS-CoV-2) variant infection has remained poorly defined. To shed light on the interaction between organs and specific SARS-CoV-2 variants, we sought to characterize the complex relationship among acute multisystem manifestations, dysbiosis of the gut microbiota, and the resulting implications for SARS-CoV-2 variant-specific immunopathogenesis in the Golden Syrian Hamster (GSH) model using multi-omics approaches. Our investigation revealed increased viremia in diverse tissues of delta-infected GSH compared to the omicron variant. Multi-omics analyses uncovered distinctive metabolic responses between the delta and omicron variants, with the former demonstrating dysregulation in synaptic transmission proteins associated with neurocognitive disorders. Additionally, delta-infected GSH exhibited an altered fecal microbiota composition, marked by increased inflammation-associated taxa and reduced commensal bacteria compared to the omicron variant. These findings underscore the SARS-CoV-2-mediated tissue insult, characterized by modified host metabolites, neurological protein dysregulation, and gut dysbiosis, highlighting the compromised gut-lung-brain axis during acute infection.

**Teaser:** In hamsters at acute infection, SARS-CoV-2 variant-specific metabolic responses and gut dysbiosis dysregulate synaptic transmission proteins.

## Introduction

The global pandemic of coronavirus disease 2019 (COVID-19) caused by the emergence of the severe acute respiratory syndrome coronavirus 2 (SARS-CoV-2) has resulted in unprecedented illnesses and deaths (*1, 2*). As of February 2024, there have been more than 774 million confirmed COVID-19 cases globally, with an estimated 7.02 million deaths (*3*). COVID-19 exhibits significant variability in clinical symptoms among individuals, spanning from asymptomatic or mild flu-like symptoms to severe pneumonia, acute respiratory distress syndrome (ARDS), multiple organ failure, and death (*1, 2*). Strain-specific immune responses and variations in disease severity were noted with the delta (B.1.617.2) variant, tending to more severe illness and pathogenicity. In contrast, the omicron (B.1.1.529) variant demonstrated increased transmissibility but less severe disease than delta (*4, 5*).

Various animal models have been used to study SARS-CoV-2 transmission and pathogenesis, including nonhuman primates and small animals (*6, 7*). Golden Syrian hamsters (GSH) infected with SARS-CoV-2 exhibit COVID-19 pathophysiology observed in humans, making them the preferred model for understanding SARS-CoV-2 disease pathogenesis and development of antiviral drugs (*8–12*). Several studies, including ours (*13*), reported comparative pathogenesis of delta and omicron in the lung and systemic inflammation (*14–17*).

While SARS-CoV-2 initially targets the respiratory system, its presence and effects extend beyond the lungs. The virus enters host cells via the interaction of its spike glycoproteins with ACE2, a receptor expressed in various cell types and tissues (*18–20*). Autopsy studies of COVID-19 patients reported the presence of SARS-CoV-2 genomic RNA or proteins in multiple organs, including lungs, heart, brain, kidney, intestine, and blood, underscoring the multi-organ involvement in COVID-19 pathology (*21–23*). No study has reported SARS-CoV-2 mediated altered metabolic regulations in distant organs despite identifying the presence of viral RNA/proteins (*24*). Viruses hijack the host’s cellular pathways for their replication, causing metabolic alterations that can significantly contribute to disease pathogenesis (*25*). Our earlier study demonstrated a distinct immunometabolic response in the lung linked to inflammation and tissue damage in the GSH model during acute infection by the delta and omicron variants, underscoring the importance of understanding the variant-specific alterations in metabolic regulations in diverse organs impacted by COVID-19 (*13*).

Respiratory viral infections, including SARS-CoV-2, can impact the gut microbiota, causing or exacerbating gastrointestinal symptoms (*26*). The presence of viral RNA in fecal samples, particularly during severe COVID-19, is often associated with gut microbiota alternations characterized by a depletion of commensal bacteria and an enrichment of pathogenic species (*27–31*). This gut dysbiosis can lead to increased intestinal permeability and the release of pro-inflammatory cytokines, contributing to systemic inflammation and neuroinflammation in the central nervous system through the gut-brain axis (*32*). This neuroinflammation contributes to the development of cognitive impairment, mood disorders, and neurodegenerative diseases often observed in the post-acute sequelae of COVID-19 (PASC) (*33–35*). Furthermore, the gut microbiota produces neurotransmitters and metabolites that can influence brain function and behavior (*36*). The impact of SARS-CoV-2-mediated gut dysbiosis on neurological symptoms reported in COVID-19 patients is not fully understood.

Integrative multi-omics studies have accelerated the context-specific understanding of complex biological processes. Therefore, the present study aimed to understand the complex relationship among acute metabolic dysregulation, dysbiosis of the gut microbiota, and the resulting implications for SARS-CoV-2 variant-specific pathogenesis at multi-systemic level. We intranasally infected GSH with SARS-CoV-2 delta and omicron variants, performed the metabolomics analysis of the brain, heart, kidney, and plasma, and used lung metabolomics from our earlier study (*13*) to identify strain-specific metabolic alterations and tissue tropism. Further, we performed whole-brain proteomics to characterize the resultant brain insult by the SARS-CoV-2 variants. Finally, we analyzed fecal and small intestine microbiomes to discern the specific microbiota selection and enrichment associated with SARS-CoV-2 variants. Our study represents a comprehensive, multi-omics systems biology investigation in the GSH model, shedding light on the pivotal role of the gut-lung-brain axis in shaping the metabolic regulation-guided disease pathogenesis during acute infections with the delta and omicron variants.

## Results

### Host tissue-specific tropism and metabolic alterations in SARS-CoV-2 variants

It has been established that SARS-CoV-2 is an multi organ disorder (*37*), and to unravel the pathogenicity of delta and omicron infection in the different tissues, and we intranasally infected GSH with delta and omicron. The GSH were euthanized four days post-infection, and collected all major organs including the brain, heart, kidney, lungs, small intestine, plasma, and feces were harvested for downstream analysis (Fig 1A). Compared to omicron-infected GSH, delta-infected GSH had a significantly higher SARS-CoV-2 genomic RNA in the lung (p=0.022), heart (p=0.009), brain (p=0.019), plasma (p=0.007), and small intestine (p=0.004). However, no significant difference in viral load was observed in the kidney and feces (Fig 1B). To understand the impact of delta and omicron infection on the metabolic responses of different tissues, we performed a targeted metabolomic analysis of brain, heart, kidney, and plasma samples. Like the lung metabolic profile (*13*), a distinct clustering was observed between delta and omicron in the brain and plasma but not in the heart and kidney metabolomics (Fig 1C). The differential metabolite analysis (DMA) identified a tissue-specific metabolite pattern in amino acid and lipid metabolism (Fig 1D). In the brain, compared to uninfected controls, distinct patterns were observed with 114 and 53 differentially abundant metabolites detected in the delta and omicron-infected hamsters, respectively (adjusted p<0.1) (Fig 1D and Table S1).

**Figure 1:**
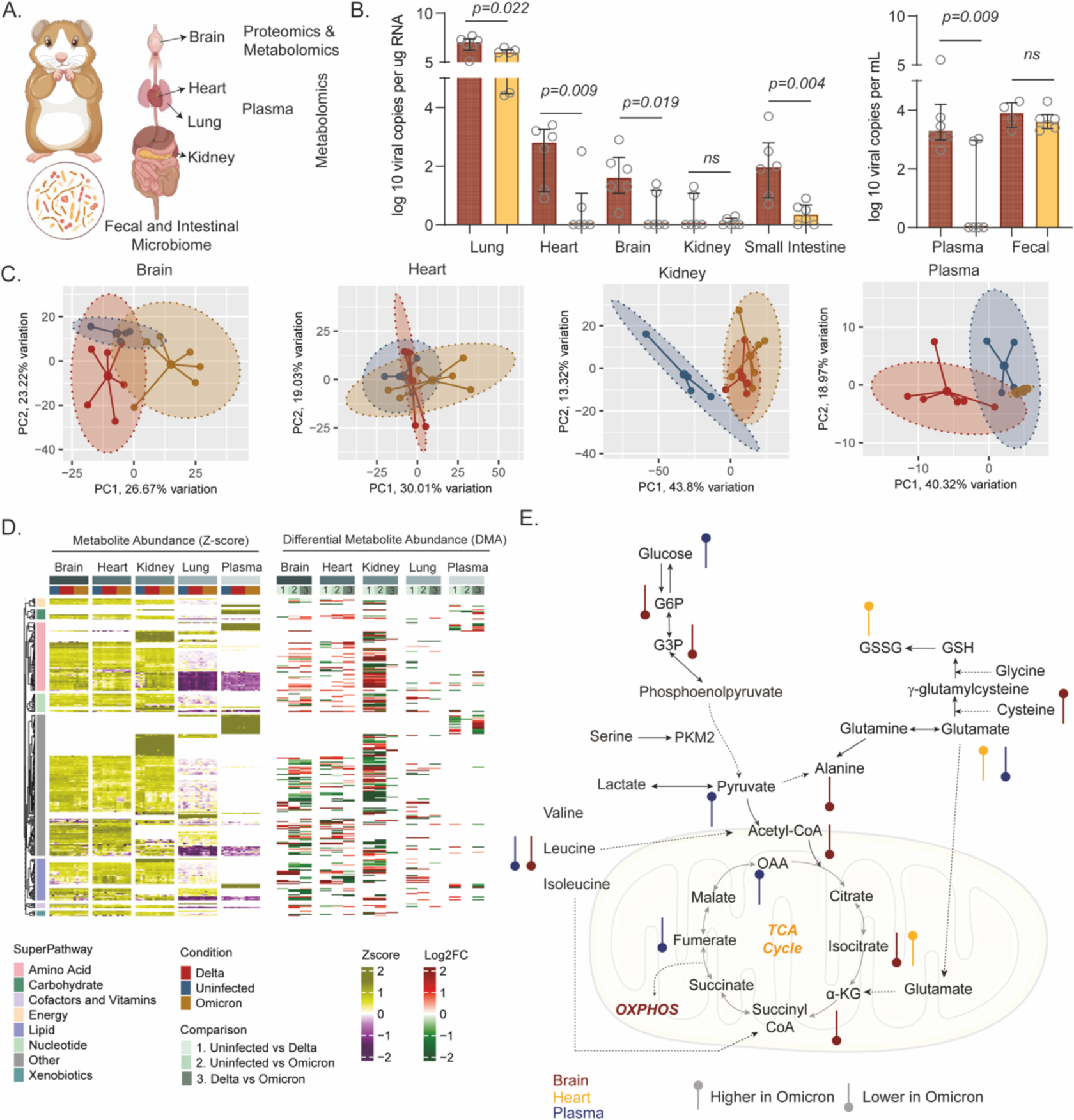
Tissue-Specific Tropism and Metabolic Changes in SARS-CoV-2 Variants. **(A)** Schematic representation of the experiment design and data acquisition. **(B)** Bar graph showing the viral load in different tissues infected with delta and omicron variants. **(C)** PCA-based sample distribution was generated using metabolomics data derived from other tissues. Log2 scaled metabolomics data was used for the visualization. The ellipse represents 90% confidence space, and the lines connect each sample with the corresponding centroid. **(D)** Heatmap visualization of metabolite abundance in each tissue. The heatmap shows the enrichment pattern of metabolites and related log2 fold changes of enrichment. Rows are metabolites significantly enriched (p.adj<0.1) in any tissues, and columns are samples. **(E)** Schematic diagram of metabolic reactions showing dysregulation of key metabolites in omicron variant infected tissues compared to delta variant infected tissues.

Interestingly, metabolites related to central carbon metabolism, glucose-6-phosphate, alpha-ketoglutarate, and isocitrate, including nicotinamide adenine dinucleotide phosphate (NADP+), were downregulated in omicron infected brain (Fig 1E). This pattern suggests that SARS-CoV-2 variants result in differential metabolic perturbations related to energy metabolism that could disrupt the balance of cellular redox homeostasis, potentially leading to oxidative stress and enhanced mitochondrial dysfunction. In contrast, the alterations of the metabolites were mainly observed between the uninfected and SARS-CoV-2 variants in heart and kidney but not between the variants [65 and 16 altered metabolites in heart (Table S2) and kidney (Table S3) respectively]. At the systemic circulations (in plasma), omicron-infected GSH exhibited fewer metabolite alterations than controls (Fig 1D). In contrast, delta infection had a more significant impact, with 37 significantly altered plasma metabolites when compared to controls and 48 when compared to omicron-infected GSH (Table S4). Strikingly, plasma metabolomics identified a specific pattern with increased glycolysis components, e.g., glucose, pyruvate, and oxaloacetate, but decreased glutamate and fumarate (Fig 1E) in omicron, indicating a shift in energy metabolism to increased glucose utilization and glycolytic activity, which are often associated with cellular stress and inflammation (*38, 39*).

### Dysregulation of post-synaptic proteins in delta-infected GSH

We observed an increased level of glutamate in circulation and severe depletion of essential and non-essential amino acids in the brain of omicron, which may impact neurotransmission and neuropathogenesis associated with alterations in amino acid metabolism. Therefore, we performed a quantitative proteomic analysis of brain tissue obtained from delta and omicron-infected GSHs. The PCA of the brain proteome data showed distinct clusters of delta and omicron-infected GSHs (Fig 2A). A distinct pattern of the differentially expressed proteins was observed in the delta and omicron-infected GSHs, while the uninfected GSHs were heterogeneous (Fig 2B and Table S5). We performed the directionally based gene set enrichment analysis in the KEGG database. We identified the downregulation of most metabolic pathways, including amino acid metabolism, in delta, further supporting the metabolomics profile (Fig 2C and Table S6). In line with the metabolic pathways, the synaptic processes, e.g., serotonergic synapse, dopaminergic synapse, and GABAergic synapse, were also downregulated in delta-infected GSHs along with the critical signaling pathways that regulate the metabolism, e.g., PI3K-Akt, mTOR signaling. Gamma-aminobutyric acid (GABA) was significantly low (adjusted p=0.033) in delta-infected brains, and serotonin was lower in delta than omicron. However, these differences did not reach statistical significance (Fig 2D). Identifying protein commonalities among synaptic, metabolic, and signaling processes reveals an overlap in PI3K-Akt and mTOR signaling pathways (Fig 2E).

**Figure 2:**
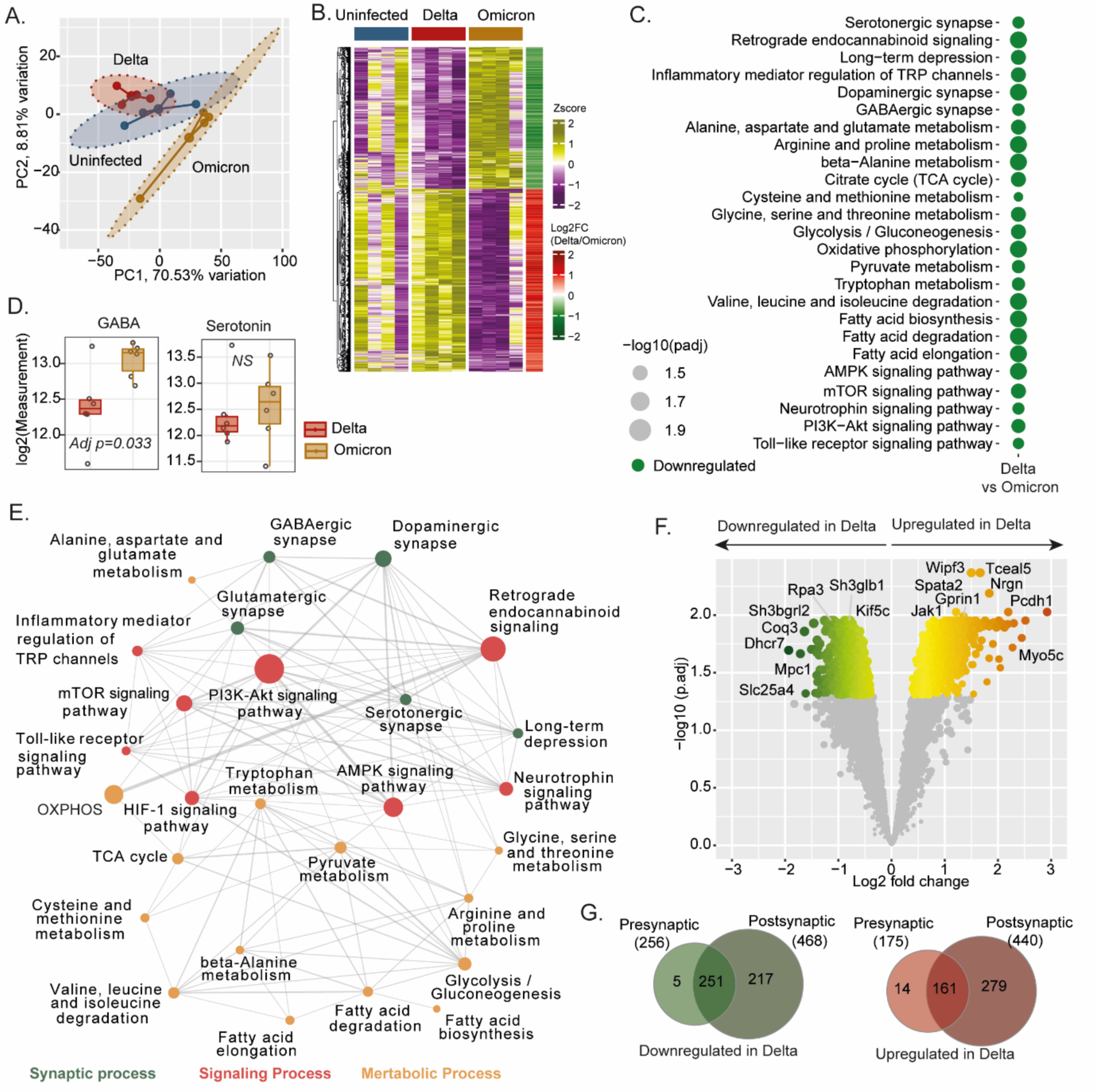
Proteomics analysis of the SARS-CoV-2 variant-infected GSH. **(A)** PCA-based sample distribution was generated using brain proteomics data. Quantile normalized proteomics data was used for the visualization. The ellipse represents 90% confidence space, and the lines connect each sample with the corresponding centroid. **(B)** Heatmap visualizing protein expression pattern in uninfected, delta, and omicron infected brain tissue and expression pattern of significantly regulated (p.adj < 0.05) protein in delta infected brain compared to omicron infected brain samples. Rows are proteins, and columns are samples. **(C)** Bubble plot showing pathway enrichment analysis results. The plot indicates significantly enriched (p.adj < 0.05) neuronal, metabolic, and metabolism-associated pathways in delta compared to omicron-infected brain samples. Bubble size is relative to negative log 10 scaled adjusted p.value of enrichment. **(D)** Box plots showing the level of GABA and serotonin in the brain tissue of delta and omicron-infected hamsters. **(E)** Network visualization of significantly enriched pathways shown in 2C. Edges represent protein overlap between the corresponding pair of pathways. Edge width is relative to the number of proteins overlapping between the paths. Node size is relative to the number of significantly regulated proteins in the corresponding pathways. **(F)** Volcano plot visualizing differential expression analysis results of pre-synaptic and post-synaptic proteins in delta infected brain compared to omicron infected brain. Significantly regulated (p.adj < 0.05) key synaptic proteins are labeled. **(G)** Venn diagram of pre-and post-synaptic proteins upregulated and downregulated in delta-infected brain compared to omicron-infected brain samples.

Furthermore, our proteomic analysis revealed that synaptic processes got dysregulated, and linked with amino acid metabolism (Fig 2E). Based on this observation, we posit that downregulated metabolic processes in the delta-infected GSHs may contribute to differential synaptic pathways. Therefore, we further looked for the level of synaptic proteins between the delta and omicron infected GSH. In delta-infected GSH, we observed a significant decrease in the expression of several synaptic proteins, including Rpa3, Sh3glb1, Kif5c, Dhcr7, and Mpc1. In contrast, synaptic proteins such as Wipf3, Spata2, Jak1, Nrgn, and Myo5c showed increased expression in delta compared to omicron-infected GSH (Fig 2F). Notably, 615 synaptic proteins had a significantly higher expression, of which 175 were presynaptic, and 440 were postsynaptic proteins, in delta compared to omicron. Of the 724 synaptic proteins with substantially lower expression, 256 are presynaptic, and 468 are postsynaptic proteins in delta compared to omicron. (Fig 2G). These data suggest that delta infection resulted in differential synaptic function compared with omicron infection.

### Impairment of gut homeostasis in delta-infected GSH through selective bacterial enrichment

We observed a more significant difference between variant infections in GSH in lung, brain, and plasma metabolic profiles. We hypothesized that overall synaptic transmission and plasticity changes may be mediated by systemic factors and signaling mechanisms through the gut-lung-brain axis. Therefore, we assessed the fecal microbiota composition in GSH infected with the SARS-CoV-2 variants. Shannon, Simpson, and Chao indices did not reveal significant differences in fecal microbiota diversity between SARS-CoV-2 infected and uninfected GSH (Fig 3A). However, Fisher’s alpha diversity analysis revealed a significant decrease in diversity in GSH infected with the omicron variant compared to those infected with delta at 4dpi. To comprehensively examine group distinctions, we conducted beta-diversity analyses using the Bray-Curtis index, which considers the structural composition of the microbial community. Notably, our findings showed distinct differences in the fecal microbiota composition among delta-infected, omicron-infected GSH, and the naïve control group (Fig 3B). Fecal microbiota composition at the family level did not differ significantly between the groups.

**Figure 3:**
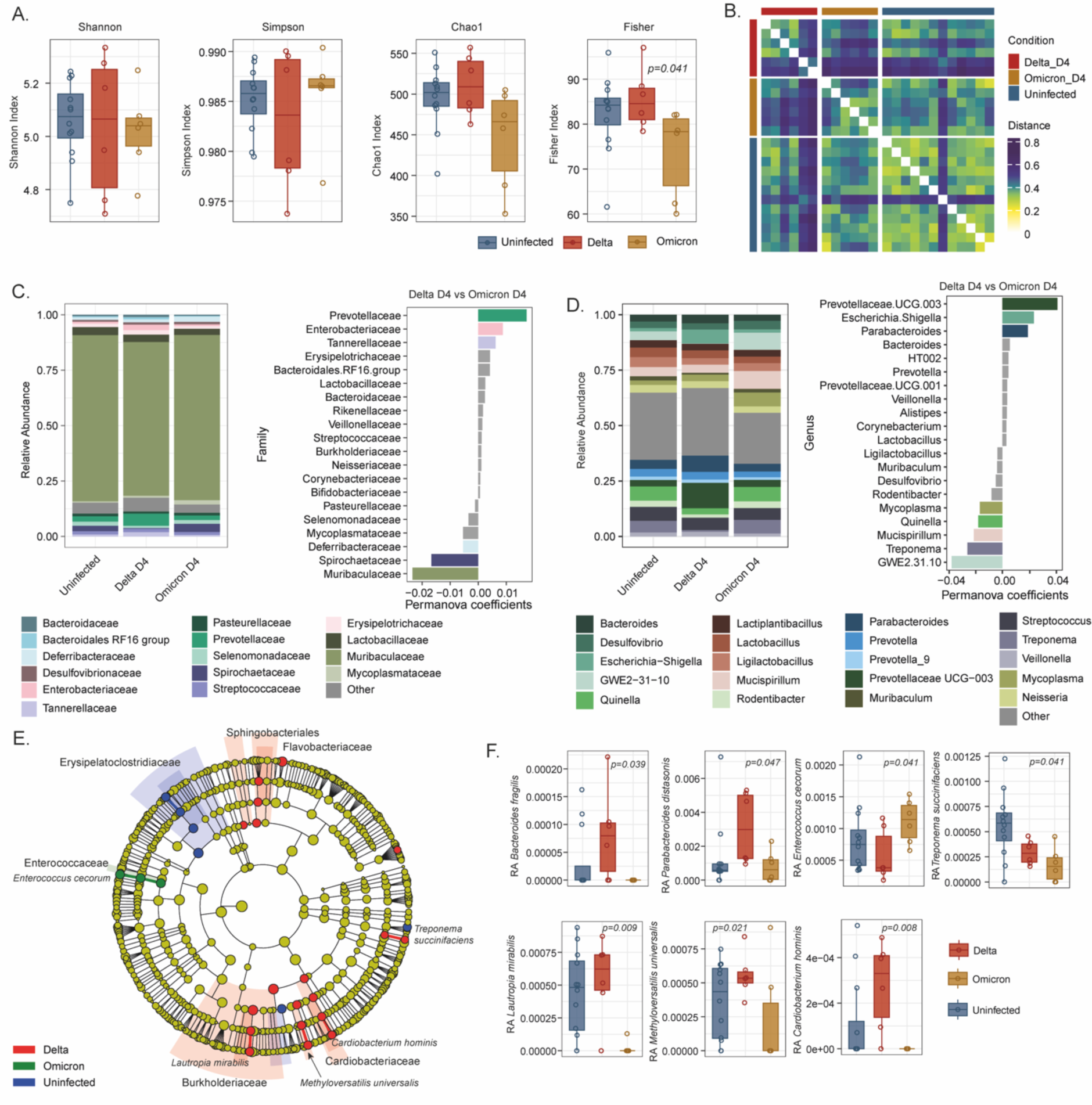
Fecal Microbiome analysis of the SARS-CoV-2 variant-infected GSH. **(A)** Box plots of various alpha diversity indices computed from fecal microbial data. The plot shows the richness and evenness of microbial diversity in the samples. **(B)** Heatmap visualizing beta diversity between the fecal samples. Distance between samples calculated using the Bray-curtis method is plotted in the heatmap. Row and column annotation represents sample conditions. **(C)** Bar graph of relative abundance and PERMANOVA analysis results at family taxa level. Average relative abundance of samples of each group is visualized in the stacked bar graph. The top 20 microbial families differentiating between delta and omicron faecal samples identified from PERMANOVA analysis are plotted in the bar graph. **(D)** Bar graph of relative abundance and PERMANOVA analysis results at genus taxa level. The average relative abundance of samples of each group is visualized in the stacked bar graph. Top 20 microbial genera differentiating between delta and omicron day 4 faecal samples identified from PERMANOVA analysis are plotted in the bar-graph. **(E)** A taxonomic cladogram showing significant changes in bacterial populations was obtained using LDA effect size (LEfSe) analysis of the 16S rRNA sequences (LDA>3.5). **(F)** Box plots showing enrichment levels of selective microbial species identified as biomarkers in E.

However, PERMANOVA analysis displayed an increased abundance of Prevotellaceae, Enterobacteriaceae, Tannerellaceae, and Erysipelotrichaceae, all associated with inflammation, and a decreased abundance of commensal bacteria Muribaculacea and Spirochateceae in delta compared to omicron-infected GSH (Fig 3C). These changes may impact metabolic functions, particularly carbohydrate metabolism and energy expenditure. At the genus level, delta-infected GSH had a higher abundance of *Prevotellaceae.UCG.003*, *Escherichia-Shigella*, *Parabacteroides* and lower abundance of *GWE2.31.10*, *Treponema*, *Mucispirillum*, *Quinella* compared to omicron-infected GSH (Fig 3D). Linear discriminant analysis effect size (LEfSe) further showed the taxonomic biomarkers in different groups. In delta-infected GSH, our analysis revealed the presence of five distinct biomarker taxa: Flavobacteriaceae, Actinomycetaceae, Rhodoycclaceae, Burkholderiaceae, and Cardiobacteriales that can impact gut homeostasis. Conversely, in omicron-infected GSH, only the biomarker taxon Enterococcaceae was detected (Fig 3E). Additionally, when we conducted individual comparisons at the species level, we found a significant increase in the abundance of *Bacteroides fragilis, Parabacteroides distasonis*, *Lautropia mirabilis, Methyloversatilis universalis, Cardiobacterium hominis* in delta-infected GSH when compared to omicron-infected GSH. Furthermore, we observed a decrease in the abundance of *Enterococcus cecorum* and *Treponema succinifaciens* (Fig 3F). Thus, our data indicate that the preferential enrichment of specific bacterial strains characterizes a disruption of gut homeostasis in GSH infected with the delta variant linked to the inflammation and metabolic profile.

### Distinct small intestine microbiota compositions in delta vs. omicron-infected GSH

We also evaluated the composition of the small intestine microbiota in GSHs infected with the delta and omicron variants, where we did not observe any significant differences in diversity between infected vs uninfected hamsters (Fig 4A). However, beta diversity analysis revealed distinct compositions of the small intestine microbiota in infected versus uninfected hamsters (Fig 4B). While the overall composition of the small intestine microbiota at the family level did not appear to differ significantly between the groups (Fig 4C), at the genus level, we noted a significant increase in the abundance of *Bifidobacterium* and *Escherichia-Shigella* in delta-infected hamsters compared to omicron-infected hamsters (Fig 4D). The co-expression analysis between the brain and plasma metabolomics, small intestine, and fecal microbiome identified a more significant association among the microbiome-associated metabolites in brain metabolomics that can explain the role of the gut-lung-brain axis. *Lautropia* was positively associated with plasma fumarate, beta-alanine, and brain methionine in delta infection, indicative of variant-specific host-pathogen interactions that influence metabolic pathways differently (Fig 4E).

**Figure 4:**
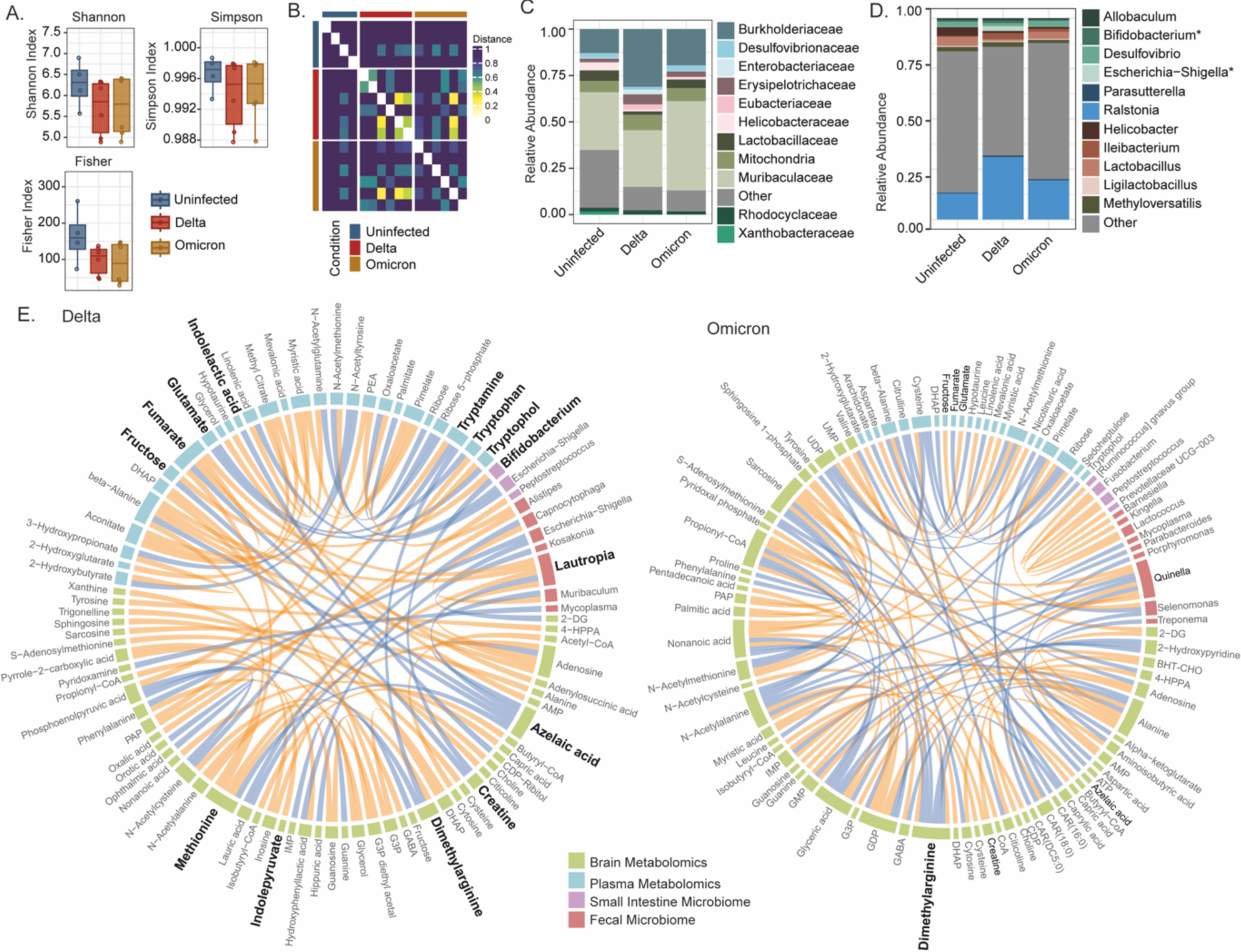
Microbiome analysis of the SARS-CoV-2 variant-infected GSH small intestine and multi-omics association. **(A)** Box plots of various alpha diversity indices computed from small intestine microbial data. The plot shows the richness and evenness of microbial diversity in the samples. **(B)** Heatmap visualizing beta-diversity between the small intestine samples. Distance between samples calculated using the Bray-Curtis method is plotted in the heatmap. Row and column annotation represents sample conditions. **(C)** Bar graph of relative abundance at family taxa level. The average relative abundance of samples of each group is visualized in the stacked bar graph. **(D)** The bar graph of relative abundance at the genus taxa level. The average relative abundance of samples of each group is visualized in the stacked bar graph. Asterisks denote significance (p-value < 0.05) by the Mann-Whitney U test between delta and omicron-infected samples. **(E)** Chord-diagram visualizing association between brain and plasma metabolomics and small intestine and fecal microbiome in delta and omicron infected samples. The links denote significant (p.adj < 0.05) Spearman’s correlation. Significantly enriched features were used to compute the correlation. Steel blue-colored links represent negative correlations, and orange-colored links represent positive correlations. Features with many correlations (adjusted p<0.00005) are bolded.

## Discussion

Our study employed the GSH model to delve into the nuanced host tissue responses to SARS-CoV-2 infection, emphasizing the delta and omicron variants. Our data suggests that the delta variant has a more robust ability to replicate and spread in various body parts than the omicron variant, leading to more severe and widespread infection. Notably, the SARS-CoV-2 delta and omicron variants exhibit diverse tissue-specific metabolic perturbations associated with energy metabolism, posing a risk to the equilibrium of cellular redox homeostasis that can contribute to oxidative stress and mitochondrial dysfunction. Further, brain proteomics indicates dysregulation of synaptic transmission, plasticity, and imbalance of excitatory and inhibitory signals within the brain in a variant-specific manner. Finally, the co-expression analysis unveiled a more pronounced association between gut microbiome-associated metabolites in brain metabolomics, emphasizing the potential involvement of the gut-brain axis in neurological consequences of the delta variant.

Although COVID-19 was initially recognized as a lung disease, it is now known to affect extrapulmonary organs, including the brain, heart, kidney, and intestine (*21–23*). In our GSHs model, infection with omicron has shown reduced viral burden in the lungs, heart, brain, plasma, and intestine compared to the delta variant. This aligns with previous reports describing the attenuated replication kinetics of omicron in human lung tissues and ex *vivo* models (*40–43*).

While numerous studies have focused on unraveling the pathogenesis of SARS-CoV-2 and the pathophysiology of COVID-19 (*12, 44–47*), the impact of SARS-CoV-2 variants on metabolic regulations in diverse host tissues during acute infection remains elusive. Our study profiled metabolites in different tissues from GSHs infected with delta and omicron at acute infection. The distinct metabolic responses in multiple tissues and plasma of delta and omicron-infected GSHs suggest that these variants might elicit unique host responses. Omicron infection revealed dysregulation of numerous brain metabolites, specifically showing upregulation of dimethylarginine and amino acids and downregulation of α-ketoglutarate. These associations underscore the potential neurological implications of omicron infection, raising concerns about its impact on brain haemostatics (*48–50*). Furthermore, the downregulation of α-ketoglutarate, a metabolite with known neuroprotective and anti-aging properties, in omicron-infected brains suggests potential consequences for cellular metabolism and overall neurological well-being. A previous study reported an association between increased levels of symmetric dimethylarginine (SDMA) and Alzheimer’s disease (*48*). An elevated level of aspartic acid, phenylalanine, methionine, and tyrosine in the brain is linked to Alzheimer’s dementia (*48*). Additionally, a pilot study reported a higher concentration of glycine, methionine, and phenylalanine in the cerebrospinal fluid of dementia patients compared to healthy controls (*49*). There are several metabolites reported to be dysregulated in the brains of AD, Dementia, and Schizophrenia, patients including cystine, lactic acid, S-Adenosylhomocysteine, valine, lysine, leucine, and glycine, among others. These metabolites are also dysregulated in the SARS-CoV-2-infected hamster brains at acute infection. Future studies should determine if these dysregulated metabolites persist even after the recovery from active disease and if they are linked to the development of neurological symptoms reported in PASC patients.

Several previous studies assert that COVID-19 is associated with neurological symptoms (*51*). Our study analyzed the brain proteome of omicron and delta-infected GSH. Interestingly, delta infection was associated with significantly lower expression of post-synaptic proteins. Disruptions in post-synaptic protein signalling pathways have well-documented associations with cognitive impairments, synaptic dysfunction, and various neurological disorders, including Alzheimer’s disease (AD), Parkinson’s disease (PD), and schizophrenia (*52–57*). Moreover, the dysregulation of post-synaptic proteins may disrupt synaptic plasticity, impairing the brain’s ability to adapt and form new memories, a crucial aspect of cognitive function (*55*). Notably, higher levels of synaptic proteins, including GAP43 and neurogranin are detected in the spinal fluids of early-stage AD patients (*58*). Hence, the dysregulation of pre/post-synaptic proteins in the SARS-CoV-2 infected hamsters brains may be responsible for the neurological consequences of SARS-CoV-2 infection.

Further, delta-infected GSH showed increased levels of α-synuclein (SNCA) and tau (MAPT) protein in the brain. These findings parallel what happens in progressive neurodegenerative diseases like PD and AD, where neurons upregulate α-synuclein and tau as part of their respective stress responses (*59–61*). Additionally, higher glutamate and lower GABA were observed in delta-infected GSH. Excessive glutamate activity, known as excitotoxicity, can lead to neuronal damage and cell death (*62*). It can also trigger the development of PD, AD, Huntington’s disease, seizures, schizophrenia, and stroke (*62–67*). Conversely, GABA is the primary inhibitory neurotransmitter, and its decrease means there is less inhibition of neuronal activity. This can contribute to conditions like anxiety, epilepsy, and other neurological disorders (*68*). This imbalance between excitatory and inhibitory neurotransmitters can disrupt the normal functioning of neural circuits and has been implicated in different neurological and psychiatric disorders, including major depression disorder (*69*). The balance between neural excitation and inhibition is essential for proper brain function, and its disruption contributes to neuronal disorders like AD (*70, 71*). Dysregulation of the GABAergic synapse pathway and decreased levels of GABA found in this study have also been reported in Alzheimer’s brain (*72*). Overall, our findings in the SARS-CoV-2 infected hamster brains indicate the presence of multiple molecular markers associated with the development of neurodegenerative disorders. To our knowledge, this is the first study to show the neurotransmitter dysregulation in a pre-clinical animal model of SARS-CoV-2 infection. In ongoing studies, we are examining whether the proteomics changes observed at acute infection persist post-resolution of the active disease symptoms and if these changes correlate with the developments of neurocognitive impairments observed in PASC patients.

The viral respiratory infections impact the bidirectional gut-lung axis by altering the microbial composition of the gut (*26*). Dysregulation of the gut microbiota during acute viral infections, including the influenza A virus, respiratory syncytial virus, and coronaviruses, has a negative implication for the disease pathogenesis (*26*). Our study observed that the delta and omicron variants resulted in gut dysbiosis in hamsters at acute infections. Specifically, the abundance of inflammation-associated taxa in the gut increased, including Prevotellaceae, Enterobacteriaceae, Erysipelotrichaceae, and *Escherichia-Shigella*. *Prevotella*, found in higher abundance in delta-infected GSH at the genus level, is known to be associated with heightened inflammation. This genus can stimulate epithelial cells to produce proinflammatory cytokines, triggering the recruitment of neutrophils and eliciting mucosal immune responses (*73*). Further, earlier studies also showed an increased abundance of *Escherichia-Shigella* in COVID-19 patients experiencing diarrhea compared to healthy subjects (*74*). Gut Enterobacteriaceae are shown to associate with aggregation of α-synuclein in the gut and brain and motor impairment in a mice model (*75*). Several neurodegenerative conditions are associated with an increased abundance of pro-inflammatory bacteria (*76, 77*), emphasizing the connection between SARS-CoV-2 delta infection and neuroinflammation.

Furthermore, delta infection increased the abundance of *Bacteroides fragilis* and Burkholderiaceae in the gut. A study has shown that an increased abundance of Bacteroides fragilis bacteria in the gut microbiota is associated with AD, suggesting a potential role in initiating AD-like pathologies (*78*). Specific pathogens within the Burkholderiaceae family are related to respiratory conditions such as asthma, chronic obstructive pulmonary disease (COPD), and cystic fibrosis (*79, 80*). These findings affirm the role of SARS-CoV-2 variants in gut dysbiosis and the potential systemic inflammation that can trigger neurocognitive impairments.

The study has limitations that merit comments. First, our study utilizes GSH as an animal model to investigate SARS-CoV-2 infection, which is widely used to study the SARS-CoV-2 pathogenesis (*81*). Animal models play a crucial role in initial investigations; however, it is essential to consider that variations in physiology between hamsters and humans might impact the direct applicability of the findings to human infections. Second, the study euthanizes hamsters four days post-infection, providing a snapshot of the acute infection. The dynamics of SARS-CoV-2 infection may evolve, and long-term observations could reveal additional insights into the progression and resolution of infection. Third, the study’s focus on a four-day time point may not capture the full extent of viral shedding dynamics, and more extended observation periods can provide a more comprehensive understanding. Another limitation of the current study is the number of animals used and the use of only male hamsters. This is attributed to working within the ABSL-3 suite’s limited capacity and the institutional regulations limiting the number of animals used in biocontainment facilities. However, our study is the first to employ a comprehensive approach combining virological, metabolomic, proteomic, and microbiota analyses. This multi-omics strategy provides a more holistic understanding of the impact of SARS-CoV-2 variants on various aspects of host physiology. The investigation focuses on multiple tissues, including the brain, heart, kidney, lungs, small intestine, plasma, and feces. This approach identifies tissue-specific responses to SARS-CoV-2 infection, contributing to a nuanced understanding of the virus’s pathogenicity.

In conclusion, our study investigates the complex host responses triggered by SARS-CoV-2 variants using the GSH model. Notably, these variants induce distinctive metabolic regulations in various tissues and elicit proteomic alterations in the brain. The observed dysregulation of post-synaptic proteins, coupled with the presence of markers linked to neurodegenerative disorders, underscores the potential neurological repercussions of COVID-19. Additionally, our investigation sheds light on the relevance of the gut-lung-brain axis in neurotropic viral infections, hinting at gut dysbiosis induced by SARS-CoV-2, with potential implications for neuroinflammation. Overall, our findings warrant further investigations to determine if the gut dysbiosis and metabolic and proteomic dysregulation observed at acute infection persist post-recovery from COVID-19 symptoms and their role in developing neurocognitive impairments reported in PASC patients.

## Materials and Methods

### Virus

SARS-Related Coronavirus 2, Isolate hCoV-19/USA/PHC658/2021 (Lineage B.1.617.2; Delta Variant; # NR-55611) and SARS-Related Coronavirus 2, Isolate hCoV-19/USA/GA-EHC-2811C/2021 (Lineage B.1.1.529; Omicron Variant; # NR-56481) were procured from BEI Resources. The viruses were cultured on Calu-3 cells in Eagle’s Minimum Essential Medium (ATCC 30–2003) supplemented with 10% FBS, 2 mM l-glutamine, and 100 units/ml each of penicillin and streptomycin. The viruses were tittered using Vero-E6 cells by the plaque assay described earlier(*82*). All animal experiments were performed using SARS-CoV-2 viral stock from either passage 1 or 2 in Calu-3 cells.

### Animal experiments

Eight- to 10-weeks-old male golden Syrian hamsters were acquired from Charles River Laboratories. Six hamsters per group were intranasally infected under isoflurane with 16000 PFU (100 μl) of the delta B.1.617.2 or the omicron B.1.1.529 variant. Additionally, four hamsters received PBS and served as uninfected controls. On 4-days post-infection (DPI), all hamsters were euthanized by ketamine and xylazine injection. Subsequently, 60 ml of PBS perfusion was performed, and brain, heart, kidney, lung, and small intestine tissues were harvested. Feces were collected at day 0 and 4-DPI, and blood samples were collected at 4-DPI. The UNMC IBC/IACUC approves all animal experiments in this study, and all infections with SARS-CoV-2 were carried out within an ABSL-3 facility.

### Viral load quantification

The lung, brain, heart, kidney, and small intestine tissues were homogenized in RLT buffer using a TissueLyser LT (Qiagen). RNA extraction was then performed using the RNeasy Mini Kit (QIAGEN), following the manufacturer’s specifications. Viral RNA from plasma and fecal samples was extracted from the QIAamp Viral RNA Mini Kit according to the manufacturer’s instructions. An RT-ddPCR assay was performed to quantify viral genomic (E gene) RNA in feces, plasma, and tissues. The 1-Step RT-ddPCR Advanced kit for probes, specific primer probes, and the Bio-Rad QX200 AutoDG digital droplet PCR system were utilized. SARS-CoV-2 E gene-specific primers and probe are as follows: E_Sarbeco_F1: 5′– ACAGGTACGTTAATAGTTAATAGCGT–3′, E_Sarbeco_R2: 5′– ATATTGCAGCAGTACGCACACA–3’ and E_Sarbeco_P1: 5′– FAM–ACACTAGCCATCCTTACTGCGCTTCG-BHQ1–3′. In brief, a reaction mixture of 22 μl was prepared, comprising 5μl of 4X Supermix, 2ul of 10X reverse transcriptase enzyme, 500 nM of each primer, 250 nM of the probe, and 10 μl of RNA template. This mixture was utilized to generate droplets using a QX200 droplet generator. Subsequently, the ddPCR plate containing the emulsified samples was sealed with foil, and amplification was carried out in a C1000 Touch thermal cycler. The thermal cycling conditions were: 60 min at 48°C, 10 min at 95°C, 50 cycles of a 30 s denaturation at 95°C followed by a 55°C extension for 1 min, and a final extension at 98°C for 10 min. Following thermal cycling, ddPCR plates were transferred to the QX200 droplet reader for droplet count and fluorescence measurement. Positive droplets containing amplified products were differentiated from negative droplets lacking the target amplicon by applying a fluorescence amplitude threshold, and the absolute quantity of RNA per sample was determined using QuantaSoft software.

### Metabolomics

Metabolomic profiling was conducted at the Metabolomics Core Facility, University of Iowa, Carver College of Medicine, Iowa. The samples were lyophilized and transferred to ceramic bead tubes for homogenization in extraction solvent (18-fold (w/v)). After rotating the samples at -20°C for 1 h, they were centrifuged at 21,000 *g* for 10 min. The resulting supernatants (400 µl) were transferred to 1.5 ml Eppendorf tubes, vortexed, and dried using a speed vac apparatus. The dried extracts were reconstituted in 20 µl of acetonitrile/water (1:1 v/v), vortexed, and rotated on a rotator at -20°C overnight. The reconstituted samples were centrifuged, and the supernatant was transferred to LC-MS autosampler vials for analysis. LC-MS data acquisition was performed using a Thermo Q Exactive hybrid quadrupole Orbitrap mass spectrometer coupled with either a Vanquish Flex UHPLC system or a Vanquish Horizon UHPLC system. A Millipore SeQuant ZIC-pHILIC column (2.1 x 150 mm, 5 µm particle size) with a ZIC-pHILIC guard column (20 x 2.1 mm) was used as the LC column, with an injection volume of 2 µl. The method employed a flow rate of 0.150 ml/min, with solvent A (20 mM ammonium carbonate [(NH_4_)_2_CO_3_] and 0.1% ammonium hydroxide (v/v) [NH_4_OH] with pH ∼9.1) and solvent B (acetonitrile). The gradient started at 80% B, decreased to 20% B over 20 min, returned to 80% B in 0.5 min, and was held there for 7 min. The mass spectrometer operated in full-scan, polarity-switching mode from 1 to 20 min, with a spray voltage of 3.0 kV, a heated capillary temperature of 275°C, and a HESI probe temperature of 350°C. The sheath gas flow was set to 40 units, the auxiliary gas flow to 15 units, and the sweep gas flow to 1 unit. MS data acquisition spanned a range of *m/z* 70–1,000, with a resolution of 70,000, an AGC target of 1 × 10^6^, and a maximum injection time of 200 ms. The acquired LC-MS data were processed using Thermo Scientific TraceFinder 4.1 software, and metabolite identification was based on the in-house library of the University of Iowa Metabolomics Core facility, using standard-confirmed metabolites. Signal drift correction was performed using NOREVA, and the data were normalized to the sum of all the measured metabolite ions in each sample.

### Proteomics

Samples were thawed on ice, and cut into small pieces, and 100 µL of RIPA buffer with 1 µL of 100x protease inhibitor cocktail (Roche) was added before homogenization using 400 µm LoBind silica beads (0.3-0.4 mg) and shaking on a Disruptor Genie at 2800 rpm for 2 min. This step was repeated twice; the beads were washed with 100 µL of RIPA buffer, and the supernatant was transferred to a new sample tube to precipitate proteins using chilled acetone (4x volume). The protein pellets were solubilized in 40 µL of 8M urea in 100 mM Tris-HCl, pH 8.5, and sonicated in a water bath for 10 min before 100 µL of 100 mM Tris-HCl was added. Protein concentration was determined by BCA assay (Pierce) and a volume corresponding to 25 µg of protein of each sample was taken and supplemented with Tris-HCl buffer up to 40 µL. Proteins were reduced with 0.4 µL of 0.5M dithiothreitol in Tris-HCl buffer, incubated at 37°C for 1 h and then alkylated with 0.8 µL of 0.5M iodoroacetamide at room temperature (RT) in dark for 30 min. Then 1 µg of sequencing grade modified trypsin (Promega) was added to the samples and incubated for 16 h at 37°C. The digestion was stopped with 2.6 µL cc. formic acid (FA), incubating the solutions at RT for 5 min. The sample was cleaned on a C18 Hypersep plate with 40 µL bed volume (Thermo Fisher Scientific), and dried using a vacuum concentrator (Eppendorf). Peptides, equivalent to 25 µg protein, were dissolved in 70 µL of 50 mM triethylammonium bicarbonate (TEAB), pH 7.1, and labeled with TMTpro mass tag reagent kit (Thermo Fisher Scientific), adding 100 µg reagent in 30 µL anhydrous ACN in a scrambled order and incubated at RT for 2 h. The reaction was stopped by adding hydroxylamine to a concentration of 0.5% and incubation at RT for 15 min before samples were combined and cleaned on a C-18 HyperSep plate with 40 µL bed volume. The combined TMT-labeled biological replicates were fractionated by high-pH reversed-phase after dissolving in 50 µL of 20 mM ammonium hydroxide. They were loaded onto an Acquity bridged ethyl hybrid C18 UPLC column (2.1 mm inner diameter × 150 mm, 1.7 μm particle size, Waters), and profiled with a linear gradient of 5–60% 20 mM ammonium hydroxide in ACN (pH 9.0) over 48 min, at a flow rate of 200 µL/min. The chromatographic performance was monitored with a UV detector (Ultimate 3000 UPLC, Thermo Scientific) at 214 nm. Fractions were collected at 30 s intervals into a 96-well plate and combined in 12 samples, concatenating 8-8 fractions representing peak peptide elution.

The peptide fractions in solvent A (0.1% FA in 2% ACN) were analyzed by LC-MS/MS as described before (*83*), except that mass spectra were acquired on a Q Exactive HF hybrid quadrupole Orbitrap mass spectrometer (Thermo Fisher Scientific) ranging from m/z 375 to 1700 at a resolution of R=120,000 (at m/z 200) targeting 1×10^6^ ions for maximum injection time of 80 ms, followed by data-dependent higher-energy collisional dissociation (HCD) fragmentations of precursor ions with a charge state 2+ to 7+, using 45 s dynamic exclusion. The tandem mass spectra of the top 18 precursor ions were acquired with a resolution of R=60,000, targeting 2×10^5^ ions for a maximum injection time of 54 ms, setting quadrupole isolation width to 1.4 Th and normalized collision energy to 34%.

Acquired raw data files were analyzed using Proteome Discoverer v3.0 (Thermo Fisher Scientific) with Sequest HT search engine against *Mesocricetus auratus* protein database (UniProt TaxID=10036 v2023-05-03). A maximum of two missed cleavage sites were allowed for full tryptic digestion while setting the precursor and the fragment ion mass tolerance to 10 ppm and 0.02 Da, respectively. Carbamidomethylation of cysteine was specified as a fixed modification. Oxidation on methionine, deamidation of asparagine and glutamine, and acetylation of N-termini and TMTpro were set as dynamic modifications. Initial search results were filtered with 1% FDR using Percolator node in Proteome Discoverer. Quantification was based on the reporter ion intensities.

### 16S-rDNA amplicon sequencing

Approximately 100-200 mg of fecal sample was used to isolate DNA using a spin column chromatography-based Stool DNA Isolation Kit (Norgen Biotek Corp; Product #27,600) following manufacturer recommendations. DNA from the small intestine was extracted using AllPrep DNA/RNA Mini Kit (Qiagen; catalog # 80204). DNA was quantified with a GE SimpliNano spectrophotometer and shipped on dry ice to LC Sciences, LLC (Houston, TX, USA) for 16S rRNA sequencing. A library was generated by amplifying the V3 and V4 16S rDNA gene variable regions and adding sequencing adapters and barcodes after the first cycle. Sequencing was performed with NovaSeq 6000, 2×250 bp (NovaSeq 6000 SP Reagent Kit, 500 cycles). An average of 83785 paired-end reads were obtained for each sample.

## Bioinformatics and statistical analysis

Statistical analysis of viral load between groups was performed using non-parametric Mann Whitney U test GraphPad Prism v. 10.

### Metabolomics

Principal component analysis was performed using R package PCAtools v2.6.0. Differential abundance analysis was performed using R package limma v3.50.0 (*84*). Log2 scaled metabolomics data was used for the analysis. Heatmap was created using the R package ComplexHeatmap v2.10.0 (*85*). Box plots were generated using R package ggplot2 v3.4.2.

### Proteomics

The raw proteomics data was first normalized using the R package NormalyzerDE v1.12.0 (*86*) and data normalized using the quantile method was used for all downstream analyses. Differential expression analysis was performed using R package limma v3.50.0 (*84*). Principal component analysis was performed using the R package PCAtools v2.6.0. Heatmap was created using R package ComplexHeatmap v2.10.0 (*85*). Pathway enrichment analysis was performed using R package piano v2.10.0 (*87*). The KEGG pathway gene sets obtained from enricher (*88*) libraries were used for the analysis. Venn diagram was created using the online tool InteractiVenn (*89*). Bubble, box, and volcano plots were generated using the R package ggplot2 v3.4.2.

### Microbiome

Processing raw fastq data was performed using R package dada2 v1.22.0 (*90*) and creating an amplicon sequence variant (ASV) table. Taxonomy assignment was done using Silva prokaryotic database version 138 (*91*). Downstream analysis was performed using R package phyloseq v1.38.0 (*92*). Relative abundance values were used for all the downstream analyses. Permutational multivariate analysis of variance (PERMANOVA) was done using an adonis function from the R package vegan v2.6.4. Biomarker discovery was executed utilizing the tool LEfSe v1.1.01 (*93*). Heatmap was created using the R package ComplexHeatmap v2.10.0 (*85*). Box plots and bar graphs were generated using the R package ggplot2 v3.4.2. A cladogram was created using LEfSe v1.1.01. The CHORD Diagram was created using the chordDiagram function from the R package circlize v0.4.15.

## Acknowledgment

We thank Rajesh Rajaiah and Kabita Pandey and other previous lab members of Dr. Byrareddy’s lab for technical support especially storing and procuring samples from previous study. We acknowledge the University of Nebraska Medical Center (UNMC) BSL-3 and ABSL-3 core facility for allowing us to perform all in-vivo and vitro experiments involving SARS-CoV-2. The UNMC BSL-3 core facility is administered through the Office of the Vice-Chancellor for Research and is supported by the Nebraska Research Initiative (NRI). Additionally, we would like to thank the biosafety leadership team and technical staff from the Comparative Medicine Department at UNMC for their invaluable assistance throughout the experimental procedures. We are grateful to BEI Resources for providing us with the SARS-CoV-2 virus and its variants. We are also grateful to the Metabolomics Core Facility at the University of Iowa, Carver College of Medicine, Iowa, for their expertise and support in conducting the metabolomic profile analysis.

## Author contributions

Conceptualization: SNB, UN, AA

Methodology: ATA, UKS, AA, UN, SNB

Investigation: UKS, SDJ, AV

Visualization: ATA, UN

Supervision: UN, SNB

Writing—original draft: UKS

Writing—review & editing: UKS, ATA, AA, SDJ, SNA, AV, UN, SNB

## Data and materials availability

“All data are available in the main text or the supplementary materials.” All the codes are available at GitHub:https://github.com/neogilab/CCHF_GSMM. The mass spectrometry proteomics data have been deposited to the ProteomeXchange Consortium (http://proteomecentral.proteomexchange.org) via the PRIDE partner repository with the dataset identifier PXD048058 (*94*). Metabolomics data have been deposited in the figshare: 10.6084/m9.figshare.24812082. 16S rRNA sequencing files have been deposited in the NCBI Sequence Read Archive with the Bioproject ID: PRJNA1060325.

